# Community-level plant functional strategies explain ecosystem carbon storage across a tropical elevational gradient

**DOI:** 10.1101/2025.02.08.637226

**Authors:** Dickson Mauki, Peter Manning, Matthias Schleuning, Andreas Hemp, David Schellenberger Costa, Natalia Sierra Cornejo, Joscha N. Becker, Andreas Ensslin, Gemma Rutten, Margot Neyret

## Abstract

1. Plant functional traits play an important role in shaping plant ecological responses to environmental conditions and influencing ecosystem functioning. However, how whole-plant functional strategies manifest at the community level to influence aboveground and belowground carbon storage across environmental gradients remains poorly understood.

2. We measured aboveground and belowground carbon stocks and the variation in whole-plant (above- and belowground) functional strategies at the community level in twelve ecosystem types across a broad savanna-forest-alpine elevational gradient of climate and land use on Mt. Kilimanjaro, Tanzania. Using Structural Equation Models, we disentangled the direct and land use mediated influences of climate on carbon storage from indirect influences mediated by variation in plant functional strategies.

3. We found strong coordination between above- and belowground functional traits at the whole community level, which corresponded with functional strategies related to two major trade-offs: a *slow*-conservation to *fast*-resource acquisition axis represented by a spectrum from high leaf dry matter content to high fine root nitrogen concentration; and a size-related *woody* to *grassy* community axis represented by a spectrum spanning high canopy height to high specific root length. The *slow-fast* and *woody-grassy* strategy axes were primarily driven by precipitation and land-use intensity, respectively.

4. Both functional strategies mediated the effects of climate on carbon storage. The *slow-fast* strategy axis was strongly and positively associated with aboveground carbon stocks. Meanwhile, the *woody-grassy* strategy axis was negatively associated with both aboveground carbon stocks and soil organic carbon stocks.

5. *Synthesis*. We demonstrated that major plant functional strategies manifest at the community level along elevational gradients. These strategies also explain variation in carbon storage, although aboveground storage is mostly driven by trait effects, and belowground storage by direct effects of climate. Together, these results underscore the importance of incorporating functional community data into future analysis of climate change impacts on carbon storage, which would enhance our ability to predict potential shifts in ecosystem functioning.

## 1. NTRODUCTION

Plant species differ in their allocation of resources to growth, reproduction, and survival, resulting in trait combinations that represent adaptation strategies to various environmental conditions (Chapin et al., 1993; Reich, 2014; Wright et al., 2004). Global variation in aboveground traits is described in the *global spectrum of form and function*, which differentiates two principal axes of trait variation, the first contrasting *fast*, resource-acquisitive strategies and *slow* resource-conserving strategies (Wright et al., 2004), and the second related to plant and seed size (Reich 2014; Díaz et al., 2016). Additional principal strategy axes have also been described for belowground traits, with the *collaboration* axis that describes a gradient of strategies for acquiring soil resources. This ranges from plants with a *do-it-yourself* (DIY) strategy, that acquire resources via direct root uptake to species that outsource resource acquisition by forming relationships with mycorrhizal fungi (Bergmann et al., 2020; Weigelt et al., 2021).

The major axes of functional strategy variation, initially described for individual species, can also be found manifested at the community-level due to habitat filtering selecting for consistent traits across species (Bruelheide et al., 2018). These community level trait responses are often described using abundance-weighted mean trait values (CWMs). CWMs have been shown to vary in response to environmental gradients across scales. For instance, at a global level, CWMs of plant traits related to a conservative strategy are closely correlated with precipitation gradients, while traits associated with plant size and nutrient acquisition are strongly linked to temperature (Moles et al. 2014; Bouchard et al., 2024; Joswig et al., 2022; Xu et al., 2023). Additionally, at the regional scale Li et al. (2017) observed strong associations between precipitation and CWM root traits related to mycorrhizal colonization whereas at local scale Schellenberger Costa et al. (2017) observed a positive relationship between resource acquisitive traits and precipitation.

As plant traits can affect the rate of biogeochemical processes (Chacón-Labella et al., 2023), both climate and variation in community-level strategies can influence ecosystem functioning (Freschet et al., 2010; Neyret et al., 2024; Reich, 2014). For instance, Manning et al. (2015) found significant effects of both mean annual temperature and leaf nitrogen on soil carbon, and Huxley et al. (2023) found significant direct and indirect trait mediated effects of climate on aboveground net primary productivity. Yet, most of these studies focus only on few individual traits, often leaf traits, rather than whole-plant strategy variation, and most of these studies do not incorporate root traits (Carmona et al., 2021; Cebrián-Piqueras et al., 2021; Freschet et al., 2010). This is likely due to technical difficulties in root trait measurement and a corresponding lack of data (Iversen et al. 2017; Laliberté 2017; De Deyn, Cornelissen, and Bardgett 2008) . As a result, there are still major gaps in understanding the relative importance of direct and indirect, trait-mediated climate effects in driving ecosystem functioning, especially the roles of both above- and belowground plant functional traits.

Here, we focus on one aspect of ecosystem functioning, carbon storage, which plays a crucial role in climate change mitigation (Lewis et al., 2004; Houghton, 2007). Growing evidence indicates that community-level (CWM) trait variation influences carbon storage (Conti & Díaz, 2013; Freschet et al., 2012; Vargas-Larreta et al., 2021). Specifically, the amount of wood and its carbon content are primary drivers of both aboveground and belowground carbon storage (Conti & Díaz, 2013). Accordingly, traits linked to biomass production and increased structural investment per unit biomass are expected to directly impact carbon storage (Baker et al., 2004; Moles et al., 2009). Several studies have reported associations between CWM traits and carbon, that taller vegetation with higher leaf carbon concentration stores more carbon (Shen et al. (2019), Yang et al. (2019)). Additionally, root traits can also influence both aboveground and soil carbon storage through their contribution to the fast-slow or *collaboration* strategy axis. Root traits related to the *conservation* and *outsourcing axis* (like high dry matter content and lifespan) have been associated with high carbon storage while those related to *do it yourself* and *fast* strategy axis (like specific root length and root nitrogen concentration) have been associated with high resource turnover and low carbon storage (Lachaise et al., 2022; Wen et al., 2022). However, only a few studies have assessed root trait-carbon storage relationships, and even fewer have incorporated both aboveground and belowground traits (but see Chanteloup & Bonis, 2013; Craine et al., 2002; Orwin et al., 2010).

While many trait-carbon relationships have been described, these are not universally consistent (Feng & Dietze, 2013), this may be due to scaling issues but also because the drivers of carbon storage vary across ecosystem types, with land management, and with climatic contexts (van der Plas et al., 2020; Peters et al., 2019). In addition to well-known trait responses to climatic conditions, land-use can also drive changes in plant functional strategies (Allan et al., 2015). Intensive land use, including agricultural and grazing activities, often causes a shift towards species with rapid growth and high reproductive output at the expense of species with conservative traits and longer lifespans (Laliberté et al., 2010; Schellenberger Costa et al. 2017; Flynn et al., 2011). This shift can reduce carbon storage as these fast-growing species typically have lower wood density and shorter root lifespans, leading to faster carbon pool turnover (Garnier et al., 2004). Moreover, changes in land use can disrupt relationships between plants and soil symbionts, altering ecosystem nutrients and carbon cycling (De Deyn et al., 2008), especially in poorly represented and underestimated belowground biomass in grassy/shrubs dryland biomes (Ottavian et al., 2020. Trait-carbon relationships may also differ between biomes (De Deyn et al., 2008). While tropical ecosystems are expected to play a major role in carbon storage worldwide, evidence for trait-functioning relationships from these systems is scarce (Finegan et al., 2015). There is thus a need for greater understanding of how plant functional strategies interact with land-use intensity and climatic conditions to influence carbon storage in tropical ecosystems.

Here, we assessed variation in trait composition at the community level and its relationship with above- and belowground organic carbon storage along the large tropical elevational gradient of Mount Kilimanjaro, which ranges from savanna through forest and to the alpine ecosystems. This gradient also encompasses variations in land use, as ecosystems at low to mid-elevation have experienced habitat disturbances and land use transformation (Misana, 2012). We hypothesized that the coordination of belowground and aboveground plant traits at the community level along the climatic gradients of Kilimanjaro matches the primary axes of *acquisitive* strategies to be found in ecosystems with high precipitation, high land-use intensity and moderate temperature while communities with *conservative* strategies would be found in ecosystems with low precipitation, and more extreme temperatures (Hypothesis 1).

Furthermore, we investigated whether the interplay between climate variables and plant functional strategies were related to patterns of ecosystem carbon storage. Here we hypothesised that climate influences carbon storage mostly via indirect pathways, mediated by land use and trait effects, with the highest carbon storage being in communities of tall, conservative plants (Hypothesis 2). We focus on the combined effects of climate and traits as land use effects on traits in Kilimanjaro have been described previously (Schellenberger Costa et al., 2017) though we incorporate land use effects as a control variable (Quétier et al., 2007). The hypotheses were tested using vegetation and soil data from 60 plots located along the elevational gradients of Mount Kilimanjaro. First, we assessed community level functional trait covariation using multivariate analyses Then, using Structural Equation Models, we related functional strategies to carbon stocks to quantify the relative importance of direct and indirect trait-mediated climatic influences (see Figure 1 for model structure and justification).

**Figure 1:**
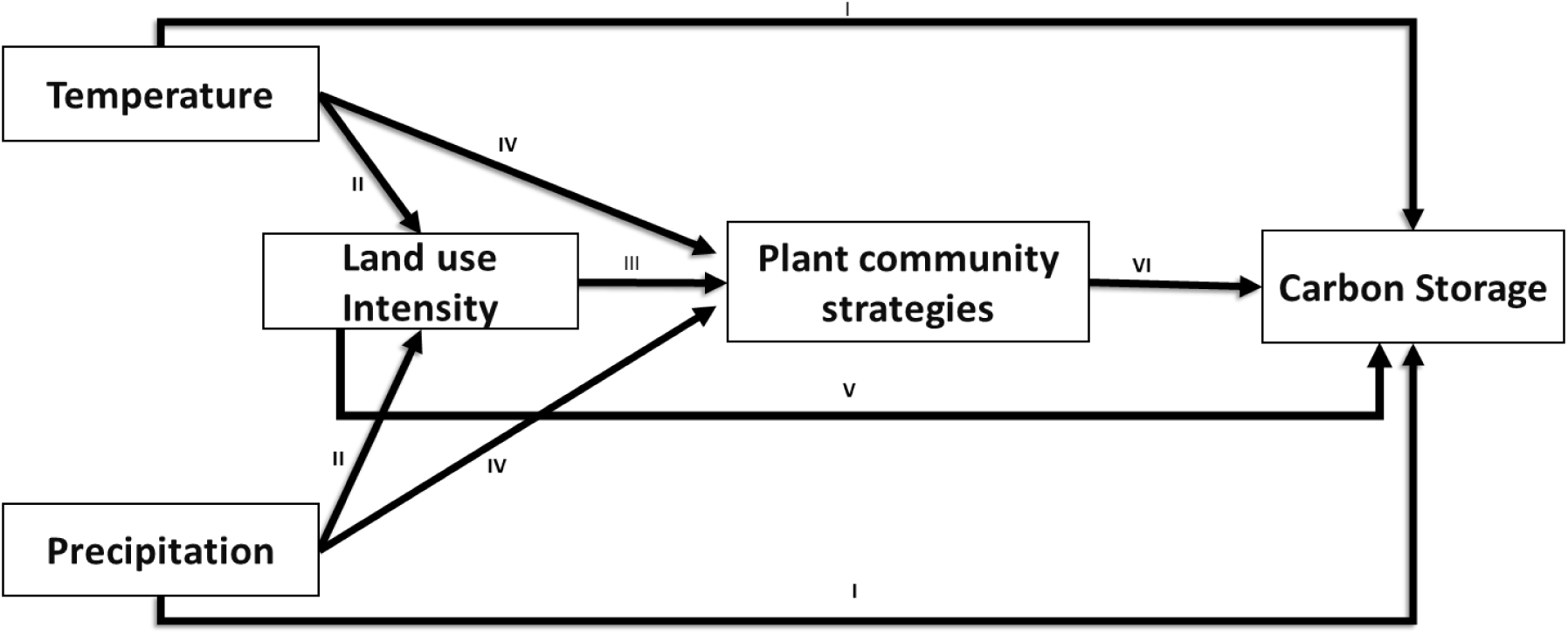
Conceptual model illustrating the plant functional strategies-carbon relationship. Arrows marked I represent direct influences of local climate on carbon storage, arrows marked II represent direct effects of climate on land-use intensity as on Kilimanjaro intensive land uses are only located in low-to mid-elevation (i.e. warmer) plots (Alemayehu et al., 2023). Arrow III represents indirect effects of climate on plant functional strategies via land use (Stampfli et al., 2018), arrow IV shows direct of influence of climate on plant strategies (Shen et al., 2016), arrow V represents the direct effect of land use on carbon storage (Ensslin et al., 2015), and arrow VI represents the influence of functional strategies on carbon storage (Chave et al., 2009).

## 2. MATERIAL AND METHODS

### 2.1 Study site

This study was conducted at Mount Kilimanjaro, northern Tanzania (3°4′33″S, 37°21′12″E) as part of the KiLi and KiLi-SES projects (Peters et al., 2019). The region is characterized by a bimodal rainfall regime with a long rain season from March to May and a short one around November (Hemp, 2006a). All soils of the Kilimanjaro massif originated from volcanic rock deposits and are therefore of similar age (Dawson 1992). The main types are andosols with folic, histic or umbric topsoil horizons, having relatively high carbon content in the upper horizons (Zech, 2006).

Field surveys were conducted in permanent plots covering 12 major ecosystem types occurring on the southern slopes of the mountain, ranging from savanna at 866 m a.s.l. to alpine vegetation at 4550 m a.s.l. (Hemp, 2006a) (Table 1). Mean annual precipitation (MAP) in the investigated sites ranges from 609 mm to 2653 mm per year along the elevation gradient, while mean annual temperature (MAT) ranges from 3^◦^C to 25^◦^C (Hemp, 2006b; Appelhans et al., 2015). A maximum of five plots were surveyed in each ecosystem type. The average distance between plots within the same ecosystem type is almost 14 km, while the minimum distance between plots is 0.2 km. Aboveground carbon was measured in 48 plots covering 10 ecosystems (Table 1). For the soil organic carbon, a total of 60 plots were surveyed covering all 12 ecosystems (Table 1).

**Table 1:**
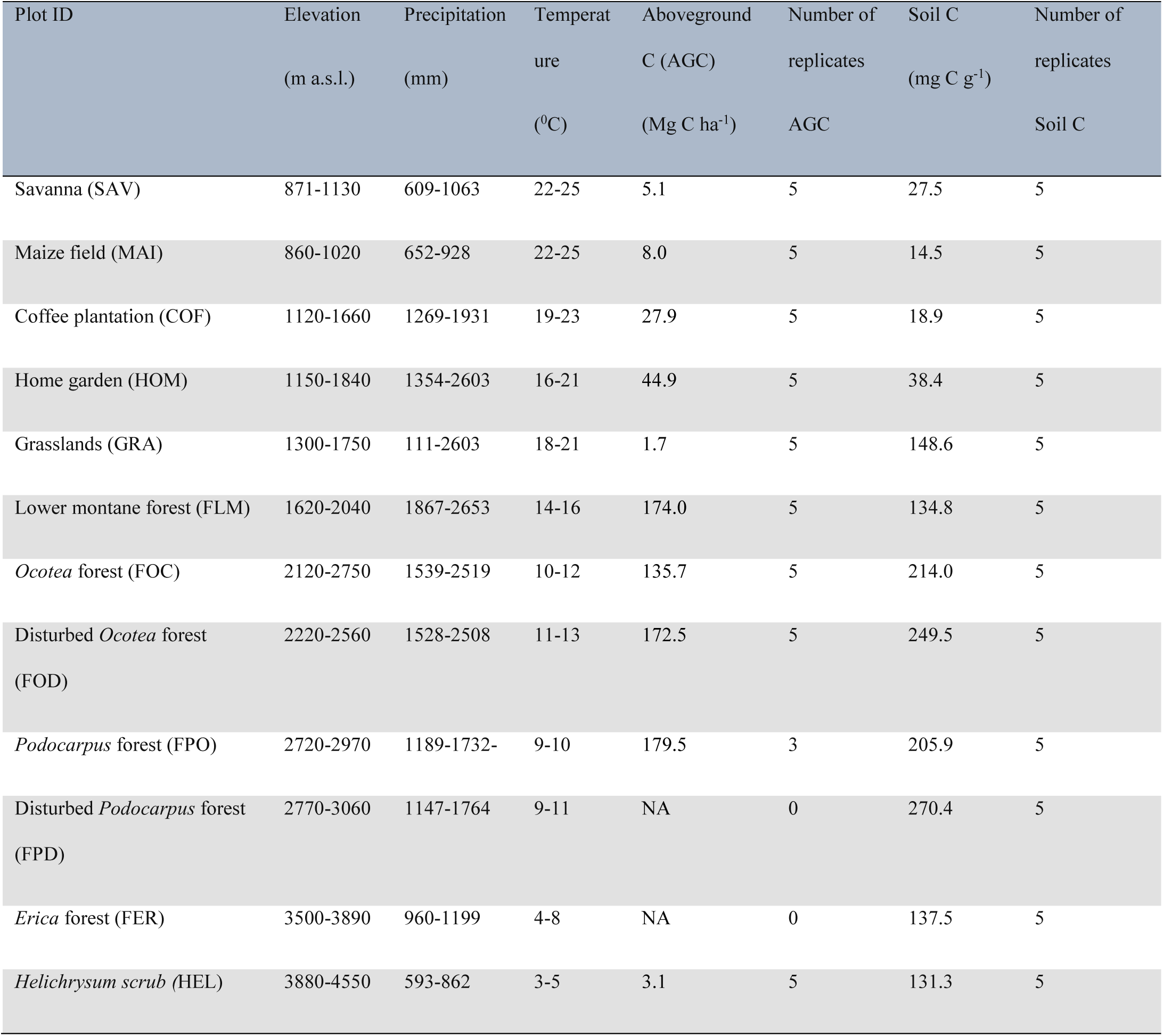
Climatic conditions, as well as measures of soil and aboveground carbon in the twelve ecosystem types surveyed at Kilimanjaro ecosystem. Elevation, precipitation and temperature values are given in range while aboveground carbon and soil organic carbon are given in mean values. NA: no measurement in the considered ecosystem.

### 2.2. Measurement of aboveground carbon storage (AGC)

To quantify AGC stocks, we used aboveground biomass data as described in Ensslin et al., (2015) including the tree, shrub and herbaceous layers. In brief, different allometric equations for different ecosystem types, and those appropriate to the vegetation type, were used to calculate the biomass in each ecosystem (Table S2), whereby the diameter at breast height (dbh) of all trees above 20 cm of DBH were measured with a diameter tape, or with a laser dendrometer for buttressed or very large trees. Tree height (H) was measured by an ultrasonic hypsometer. Shrub DBH and height was measured in subplots (5 m x 20 m) within each large plot using a diameter tape of caliper and a hypsometer. Shrubs, classed as all woody stems exceeding 1.3 m in height, but thinner than 10 cm dbh, were excluded from the tree inventory. For the herbaceous biomass, non-random sampling was used by setting four wooden frames of 50 cm x 50 cm (0.25 m^2^) within each plot to collect forbs, woody stems, mosses and lichens found at ground level. Herbaceous samples were oven-dried at 72°C for 72 hours and then weighed. For calculation of the plot level of wood biomass, wood cores were taken from the dominant tree and shrub species, accounting for 80% of the ground cover of all trees and shrubs in each plot, to measure wood density. Samples were taken from at least five individuals, if possible from different sites, at 1.3 m height. Samples were oven dried at 72^0^C for at least 72 hours and density measured as dry weight divided by fresh volume for field collected wood cores (Ensslin et al., 2015). This was complemented by data from the global wood density database (Chave et al., 2009) for 25% of species for which no core could be sampled. The carbon content of woody and herbaceous biomass was approximated as 48.2% of the total biomass per plot (Thomas & Martin, 2012). For details see Ensslin et al. (2015), and see Table 1 and Figure. S2.

### 2.3 Measurement of soil organic carbon (SOC) stocks

Soil organic carbon stocks were calculated based on SOC content and soil bulk density (BD) measurements. Mixed soil samples were collected from four corners of each plot at depths of 0– 10, 10-20, 20-30, and 30-50 cm using a standard soil auger. At each depth, samples were combined to form a composite sample for further analysis. Samples were sieved (<2mm), dried (72 h at 105°C), ground, and then analyzed for total carbon content using dry combustion at 950 °C (vario max CN, Elementar, Hanau, Germany). Additionally, at 38 plots a soil profile was established and undisturbed soil cores (100 cm³) were taken at the center of each depth interval. BD was determined as the dry weight per core volume, accounting for the volume of stones and gravel (>2mm), measured by water displacement at room temperature. For the remaining plots, BD was estimated using a pedotransfer function (PTF) *BD* = 1.94 − 0.71 *log*10(*SOC*) built on the measured data (R² = 0.86, p < 2*10^-16^). The soil carbon stock for each plot was taken as the sum of the carb on stocks in all four soil depths, with the ecosystem-types SOC being the average of the number of plots within the respective ecosystem. More details in Table 1 and Figure S2.

### 2.4 Environmental variables

Mean annual temperature (MAT) for each plot was measured in-situ from temperature loggers (data logger DK320, Driesen and Kern GmbH, Bad Bramstedt, Germany), while mean annual precipitation (MAP) was modeled using long-term observations based on a 15-year dataset from a network of rain gauges distributed across the entire mountain. The model was validated through cross-validation techniques, with performance metrics indicating strong predictive capability. Specifically, the Root Mean Square Error (RMSE) of 429 mm and the Coefficient of Determination (R² = 0.77) suggest that the model explains 77% of the observed variance in precipitation across Kilimanjaro (Hemp, 2006a; Appelhans et al., 2015). MAT decreases with elevation while MAP has a hump-shaped distribution (Figure S1). See more details in the supplementary materials.

### 2.5 Land-use intensity

We used a land use intensity (LUI) indicator developed by Peters et al. (2019), which included four key components: (a) The annual removal of plant biomass, calculated by averaging standardized estimates of plant biomass removal at study sites. (b) Agricultural inputs, derived from the average standardized estimates of irrigation, fertilization, and the use of insecticides, fungicides, and herbicides. (c) Structural characteristics, which describe the deviation from natural habitats, often used as indicators of land use. (d) A variable representing the landscape composition, defined as the proportion of agriculturally managed habitats within a 1,500-meter radius of the study area. See more explanation about derivation of the LUI in the supplementary material.

### 2.6 Functional trait and plant community data

#### 2.6.1 Aboveground traits

For aboveground traits, vegetation surveys were undertaken between 2010 and 2014 in 60 plots of 50 x 50m, slope-parallel (Hemp et al., 2021). We assessed plant abundance data based on the Braun Blanquet scale (Braun-Blanquet, 1964) and then converted abundance classes to corresponding midpoint percentage cover (e.g., a Braun-Blanquet of class 2 corresponds to 5-25% cover and the median value used was 15%). In-field plant functional trait measurements by Schellenberger Costa et al., (2017) followed the LEDA protocols (Kleyer et al., 2008; www.leda-traitbase.org).). We selected traits representative of the global trait spectrum of form and function (Díaz et al. 2016): Leaf dry matter content (LDMC), Stem-specific density (SSD), Canopy height (CH) and Specific Leaf area (SLA-) (Cornelissen et al. 2003). We aimed to measure community-level metrics from the most abundant species, constituting ≥80% of total plant biomass at the plot level (Cornelissen et al., 2003). Due to high plant diversity in this study area, it was not possible to reach this target with self-measured traits for all plots so we supplemented our field-collected traits with TRY data for missing trait values (Kattge et al., 2011). We used a total of 2377 plant traits values from 205 plant species, about 35.6% (846) was from TRY and 64.4% (1531) was from field measurements. We then used community weighted means (CWM) as a measure of functional composition. (Cavanaugh et al., 2014; Garnier et al., 2004)-function dbFD, R package FD).

#### 2.6.2 Belowground traits

For belowground traits, we did not use CWMs but rather average trait values directly measured from all roots found in soil samples, thus representing community-level traits (Sierra Cornejo et al., 2020). Soil samples were collected randomly using a soil corer of 3.5 cm diameter to 40 cm depth. In each plot, three composite samples, each obtained from mixing two samples from the field, were collected. Samples were stored in plastic bags at 5°C until processing. Roots were washed and all root fragments above 1 cm in length and below 2 mm diameter were selected. Roots were then weighed and analysis of morphological traits was conducted by scanning each root using a root perfection scanner (EPSON perfection V700 scanner (EPSON America Inc.). Specific root length (SRL, mm), mean fine root diameter (RD, mm) and root tissue density (RTD, g cm-³) were calculated from root biomass and scans using the WinRhizo software. Lastly, fine root nitrogen concentration (RNC) was measured with a CN elemental analyzer (Sierra Cornejo et al., 2020).

### 2.7 Statistical analyses

All analyses were performed in R version 4.3.1 (R Core Team, 2023).

To test hypothesis 1, we first performed a principal components analysis (PCA, package *FactoMineR*) on all community-level trait measures (above- and belowground) in which all traits were scaled to zero mean and unit variance. Based on the screen plot and Kaiser-Guttman criterion components with greater than 1 are retained (Silva et al. 2021), we retained the first two axes (PCA1_(eigenvalue)_ =3.77, PCA2_(eigenvalue)_ =1.99). We projected the scaled mean temperature precipitation and land-use intensity as supplementary variables on the PCA and tested their correlation with the two principal component axes using Pearson’s correlation coefficient. All traits and supplementary variables were transformed to Z-score before running the PCA.

To disentangle the effects of climate, land use and the functional composition on carbon storage (hypothesis 2), we used SEMs from the ’piecewiseSEM’ package (Lefcheck, 2016), which account for both direct and indirect relationships among variables in complex systems (Grace et al., 2012; Sonne et al., 2016). Our model structure mirrored hypothesized relationships (Figure 1), and we fitted separate SEMs for AGC and SOC both standardized to zero and unit variance. We used the first two axes from the PCA analysis as indicators of community-level composition variation. Paths that deviated from normality were assigned to generalized linear models with a Gaussian family within the ’*psem*’ function. To assess goodness of fit, we used Shipley’s test of d-separation via Fisher’s C statistic (*p*-value > 0.05, (Shipley, 2009)). We calculated the indirect effect of mean precipitation and temperature as the products of their path coefficients effect on functional axes and the functional composition axes effects on carbon storage. The overall total effect of each predictor on carbon storage was determined as the sum of the direct and indirect effects (Poorter et al., 2015).

## 3. RESULTS

### 3.1 Trait covariation and functional strategies

We found a coordinated response between the above- and the belowground traits and identified two axes of community level functional trait covariation (Figure. 2 and Figure. 3). The first axis accounted for 47.19% of the total variation and represented a *slow-fast* trait continuum (hereafter *slow-fast* strategy axis). This axis ranged from plant communities associated with slow-resource acquisition and long-term resource conservation, characterized by high values of leaf dry matter content, to acquisitive plant communities related to high resource capture and utilization efficiency, characterized by high root nitrogen concentration. The second axis accounted for 24.84% of the total variation and represented a *woody-grassy* trait continuum (hereafter *woody-grassy* strategy axis), ranging from plant communities dominated by *woody* traits species with higher canopy height and root tissue density to plant communities dominated by *grassy* traits with high specific leaf area and high specific root length at the high values of this axis, the latter trait indicating a DIY root foraging strategy (See Figure. 3 A).

**FIGURE 2:**
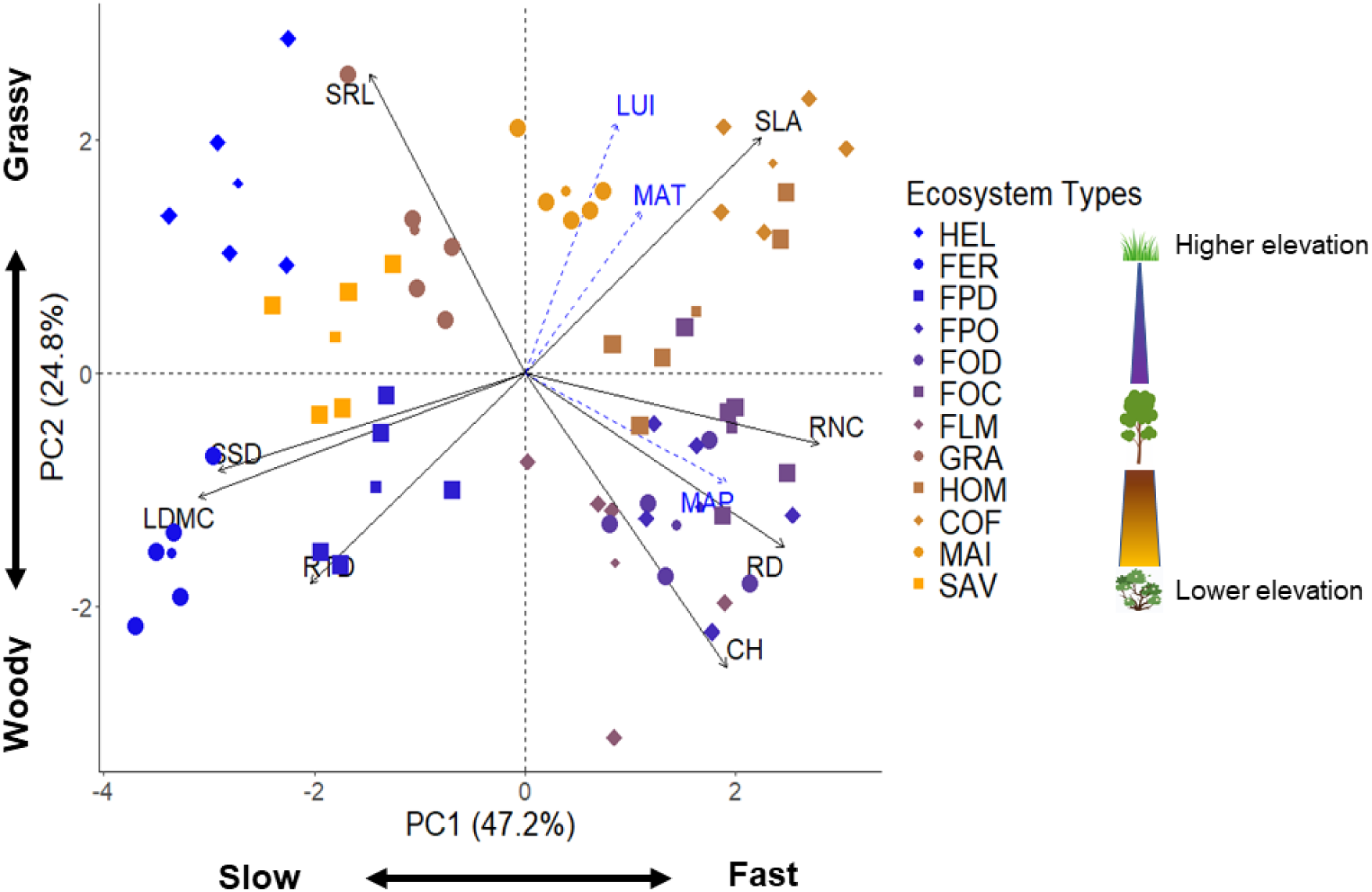
Principal component analysis of community-level below- and aboveground traits. The first axis shows the *slow-fast* strategy axis, from conservative to resource acquisitive strategies (explained variance of 47.2%). The second axis represents a transition from the woody-grassy strategy axis (explained variance of 24.8%). The abiotic drivers of the two axes are presented by the temperature (MAT), precipitation (MAP) and land use intensity (LUI) arrows, projected onto the PCA as supplementary variables. Colors indicate ecosystem types arranged according to the elevational gradients from high elevation (blue) to lower elevation (orange). The traits included are, LDMC: Leaf dry matter content, SLA: Specific leaf area, SSD: Stem specific density, CH: Canopy height, RD: Average fine root diameter, SRL: Specific root length, RNC: Fine root Nitrogen concentration, and RTD: Root tissue density. The ecosystems are; HEL-*Helichrysum*, FER-*Erica* forest, FPD-Disturbed *Podocarpus* Forest, FPO-*Podocarpus* Forest, FOD-Disturbed *Ocotea* Forest, FOC-*Ocotea* Forest, FLM-Lower Montane Forest, GRA-Grassland, HOM-Homegarden, COF-Coffee plantation, MAI-Maize fields, SAV-Savanna (Table 1).

**FIGURE 3.**
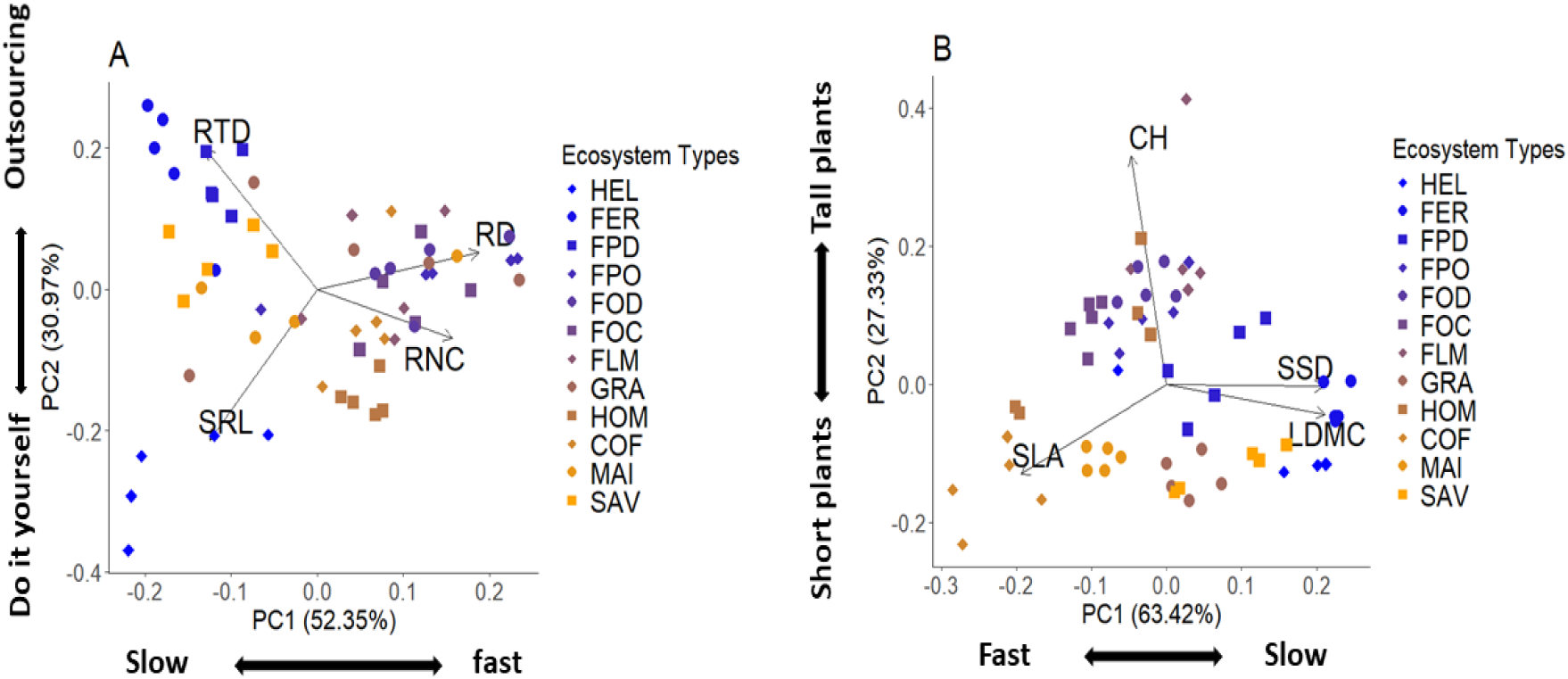
The principle component analysis shows the coordination of the A. belowground traits to form the two independent axis *slow-fast* and the *collaboration* axis. B. the aboveground traits to show the orientation of the traits to form *fast-slow* and the *plant size* axis.

Both observed functional trait axes of variation were significantly influenced by abiotic drivers. The influence of precipitation on the *slow-fast* axis (r = 0.56, *p*< 0.001) was positive and stronger than that of temperature and LUI (r = 0.31, *p*= 0.026, and r= 0.30, *p*= 0.033 respectively). For *woody-grassy* axis the influence of LUI (r = 0.54, *p*< 0.001) was positive and stronger than the positive influence from temperature and the negative influence from precipitation (r = 0.31, *p*= 0.027, and r= -0.37, *p*= 0.006 respectively, see Table S1).

### 3.2 Direct and trait-mediated effect of climate on carbon storage

Assessment of both direct and the trait-mediated influences of climate on carbon storage using structural equation models found that AGC was more strongly driven by land use- and trait-mediated influences than by direct effects of either temperature or precipitation (Figure. 4, Table 2). The SEM explained 90% of the overall variance of the direct and indirect effects of climate on AGC (Fig. 4; (χ² = 4.775, *df* = 2, p = 0.092). Specifically, the *slow-fast* strategy axis influenced AGC positively (R² = 0.33, β = 0.47, p < 0.0001), while the *woody-grassy* strategy axis influenced AGC negatively (R² = 0.59, β = -0.38, p < 0.001). Higher land use intensity was also associated with lower AGC (R² = 0.49, β = -0.33, p = 0.001). Precipitation had a direct negative relationship with the *woody-grassy* strategy axis (β = -0.47, p < 0.0001), a positive effect on the *slow-fast* strategy axis (β = 0.56, p < 0.0001), and positive relationship with AGC (β = 0.18, p = 0.007). Temperature directly and negatively influenced the *woody-grassy* strategy axis (β = -0.43, p = 0.003) and positively affected LUI (β = 0.70, p < 0.0001) (Figure. 4, Table 2).

**FIGURE 4:**
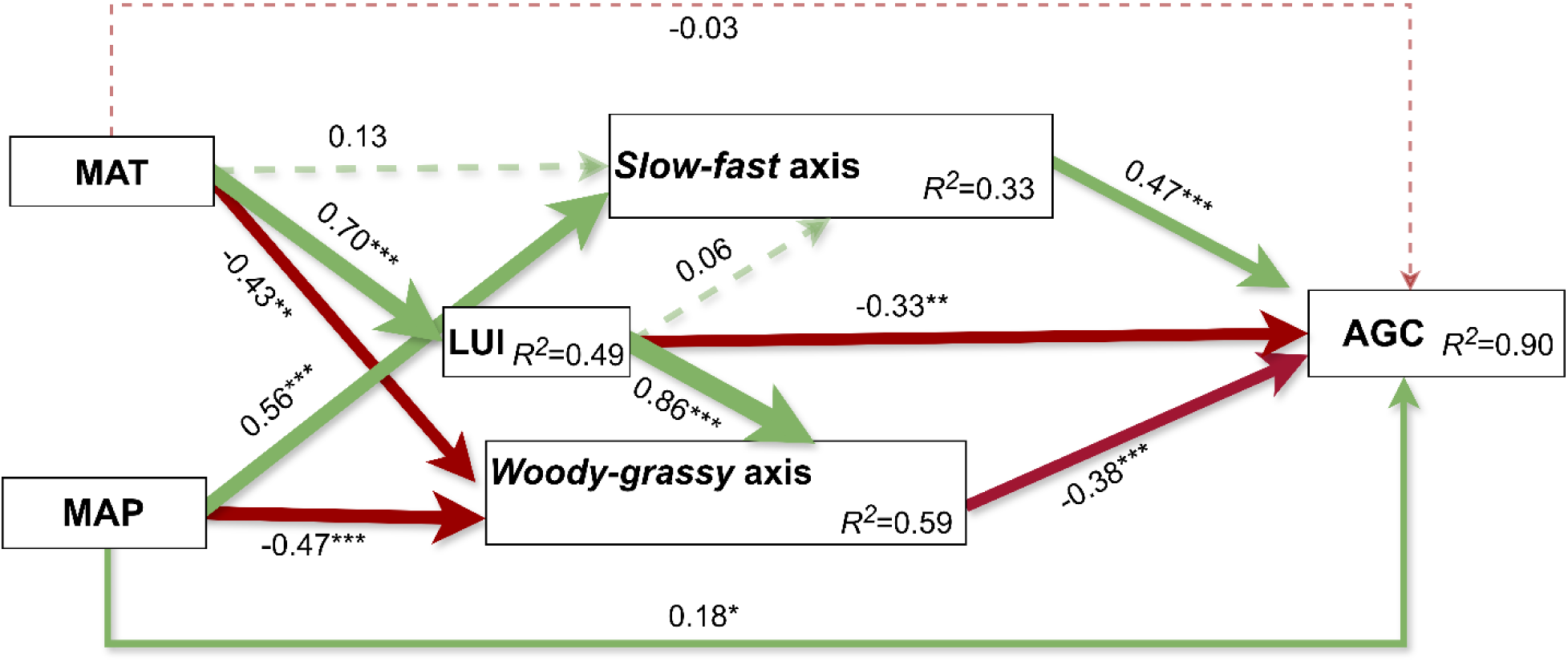
Structural equation model showing direct and indirect association of functional strategies, and their drivers, with AGC. Numbers adjacent to arrows represent the standardized path coefficients*. Slow-fast* and *woody-grassy* axes represent principal components axes of community weighted means of both belowground and aboveground plant traits variables (shown in Figure 2, Table 2). R^2^ indicates the proportion of variance explained. Solid arrows represent significant paths (\*\*\**p* < 0.001 \*\**p*<0.01 and \**p*<0.05). Dashed arrows represent nonsignificant paths. Red arrow shows negative and the green arrow shows positive paths. The arrows thickness denoted the extent of effects with thicker arrow means larger influences. Fisher’s C statistic (C = 1.11, *df*= 2, *p* = 0.574) indicated a goodness of fit (GoF) value of 0.554 for the model as well as χ² = 4.775. The Akaike Information Criterion (AIC) was 452.450, based on a sample size of N = 48 plots. MAT: mean annual temperature, MAP: mean annual precipitation.

**Table 2:**
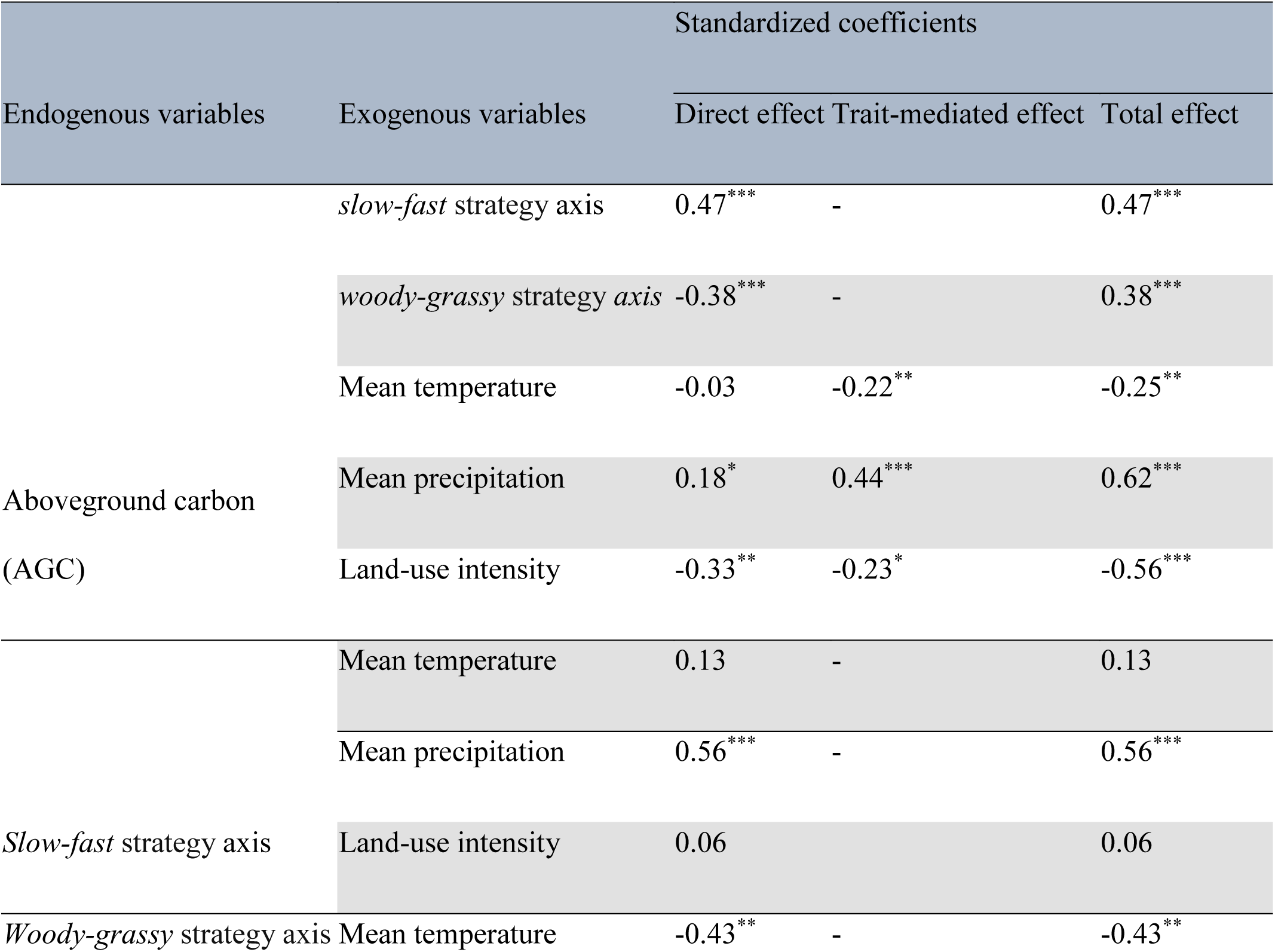
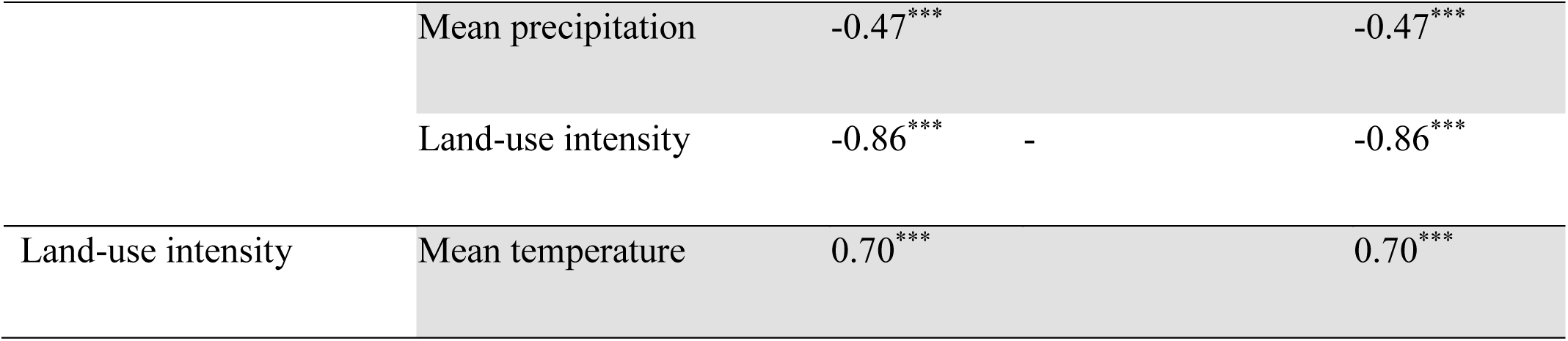
Standardized coefficients for the influence of endogenous and exogenous variables, derived from AGC structural equation model (Figure. 4). Standardized coefficients showing the direct, trait-mediated (indirect), and total effects of exogenous variables on endogenous variables. The trait-mediated effect represents the indirect influence of exogenous variables on the endogenous variable via plant functional traits (slow-fast and woody-grassy strategy axes) acting as mediators.

In contrast to AGC, SOC stocks were more strongly driven by direct than trait-mediated climate effects (Table 3). The SEM explained 63% of the overall variance of the direct and indirect effects of climate on SOC (Figure. 5; χ² = 0.155, *df*= 2, p = 0.75). Unlike the AGC model, this model had weak land use- and *slow-fast* strategy axis-mediated paths, and the *woody-grassy* axis mediated effects of precipitation were strongly, and negatively, related to SOC. (*R^2^*=0.47, β=-0.44, p < 0.0001). Precipitation had a positive direct association with the *slow-fast* strategy axis (β = 0.54, p < 0.0001) and a negative relationship with the *woody-grassy* strategy axis (β = -0.31, p = 0.003). Temperature had a positive direct association with LUI (β = 0.73, p < 0.0001) and a positive association with *slow-fast* strategy axis (β = 0.33, p =0.035). These different effects compared to what was observed in the previous model (Figure 4) are likely due to a larger sample size for SOC than AGC data, resulting in more ecosystem types being included. Finally, precipitation and temperature respectively had a positive (β = 0.21, p = 0.063, n.s). and negative (β = -0.39, p = 0.003) relationship with SOC (Figure. 5, Table 3).

**Table 3:**
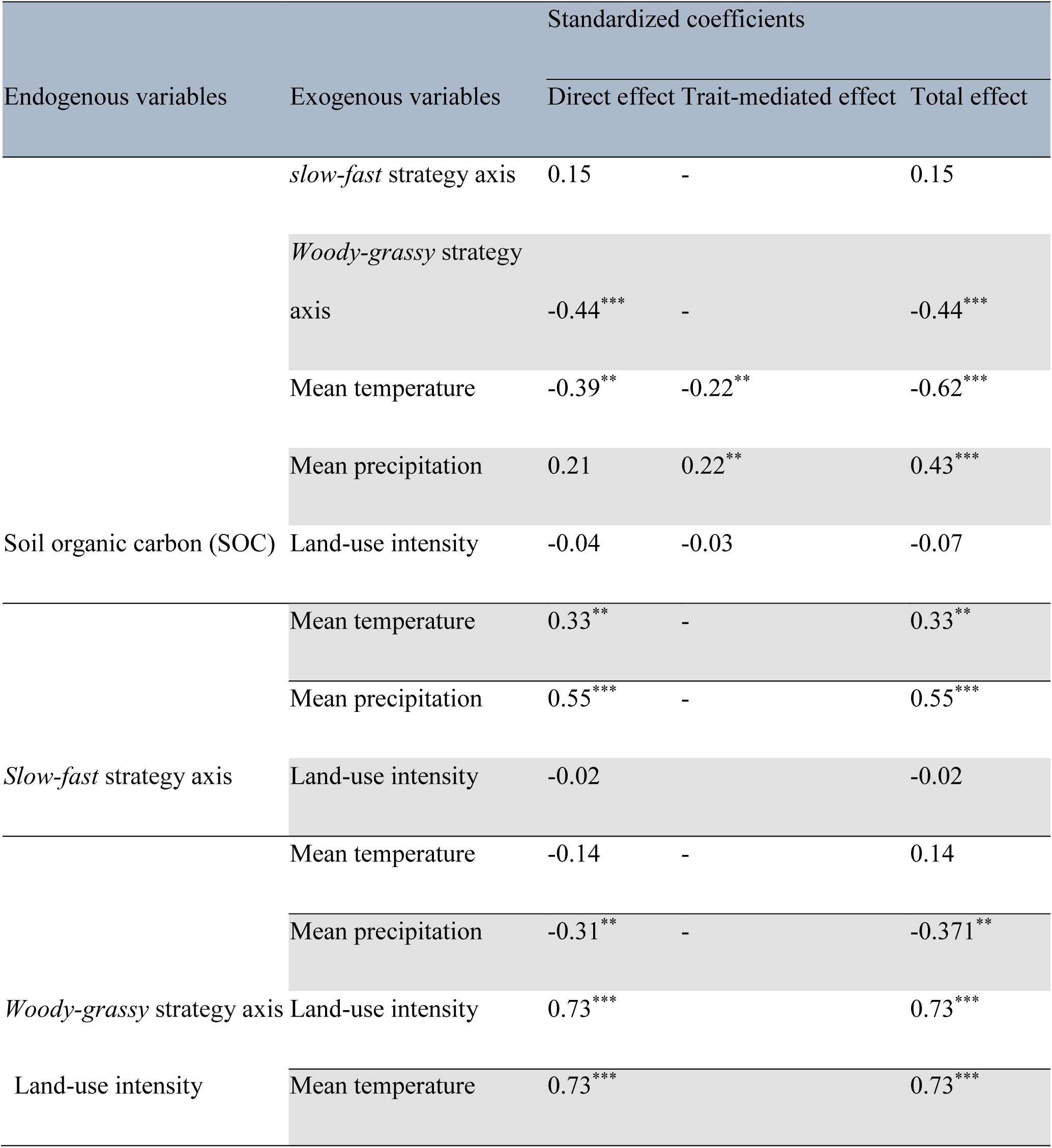
Standardized coefficients for the influence of endogenous and exogenous variables, derived from SOC structural equation model (Figure. 5). Standardized coefficients showing the direct, trait-mediated (indirect), and total effects of exogenous variables on endogenous variables. The trait-mediated effect represents the indirect influence of exogenous variables on the endogenous variable via plant functional traits (slow-fast and woody-grassy strategy axes) acting as mediators.

**FIGURE 5:**
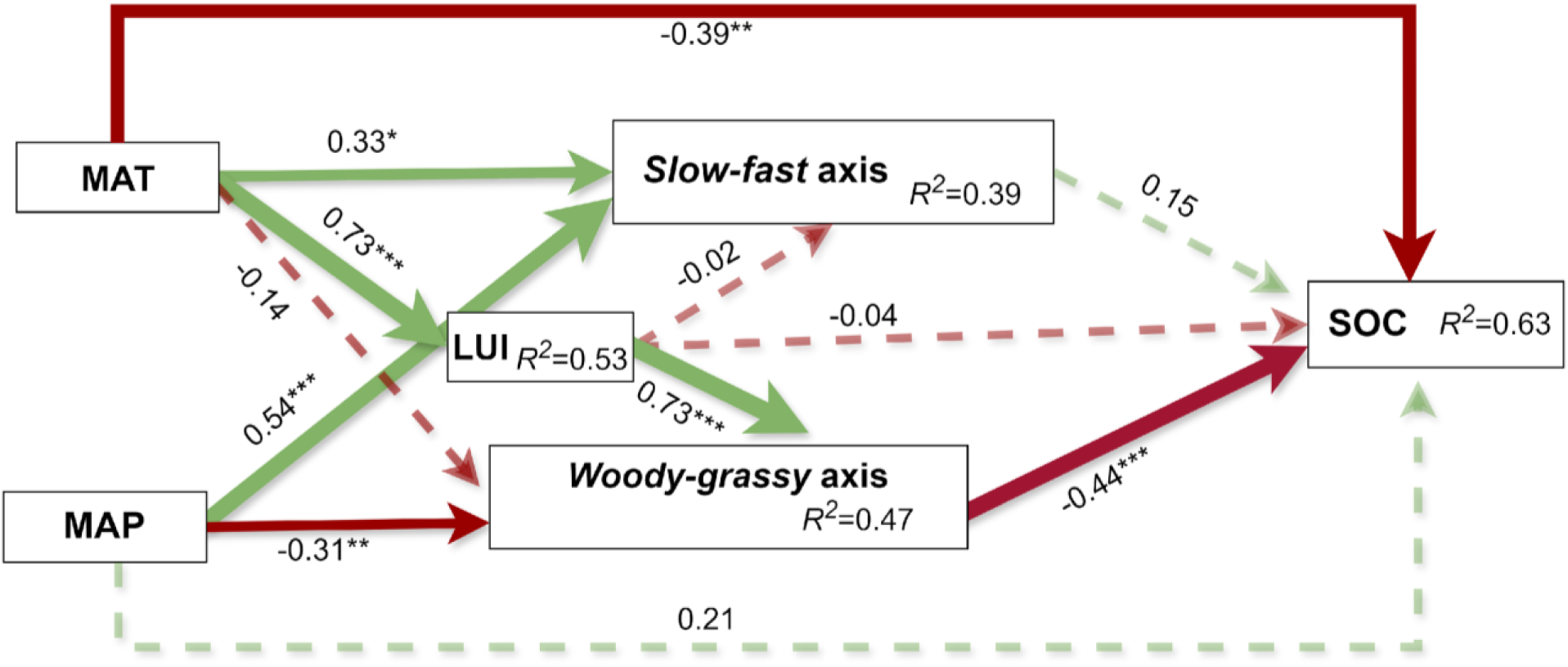
Structural equation model showing the direct and indirect effects of climate and plant functional strategies on SOC. Numbers adjacent to arrows are standardized path coefficients*. Slow-fast* and *woody-grassy* strategy axes represent principal components axes of community weighted means of belowground plant traits and aboveground plant traits. R^2^ indicates the proportion of variance explained. Solid arrows represent significant paths (\*\*\**p*< 0.001 \*\**p*<0.01 and \**p*<0.05). Dashed arrows represent nonsignificant paths. Red arrow shows negative and the green arrow shows positive paths. The arrows thickness denoted the extent of effects with thicker arrow means larger influences. Fisher’s C statistic (C = 1.965, *df* = 2, *p* = 0.75) indicated a goodness of fit (GoF) value of 0.742 for the model as well as χ² = 0.155. The Akaike Information Criterion (AIC) was 499.56, based on a sample size of N = 60 plots). MAT = mean annual temperature, MAP = mean annual precipitation.

## 4. DISCUSSION

We identified two fundamental axes of community level plant functional trait co-variation. The first, the *slow-fast* strategy axis, was mostly influenced by precipitation, and second, the *woody-grassy* strategy axis, covering both variations in root diameter and plant size, was mostly influenced by land-use intensity. Both the *slow-fast* growth and *woody-grassy* strategy axes showed strong relationships with AGC, suggesting that promoting woody and fast-growing traits in ecosystems (such as in restored forests) can maximize AGC storage and thus help mitigate climate change. However, only the *woody-grassy* axis played a crucial role in influencing SOC, which was dominated by direct climatic effects. In this study, we provide evidence that the whole-plant functional composition at the community level mediates the influence of climate. In the discussion that follows we discuss our results in more detail, starting with the axes of community level trait covariation, and then how these relate to the environment and carbon stocks.

### 4.1 Functional community strategies response to environmental gradients

By incorporating multiple aboveground and belowground plant functional traits, we identified the manifestation of whole-plant strategy axes at the community level. The first axis differentiated between *slow* and *fast* communities and captures variation in plant community strategies from low-resource, *slow*-growth conditions to high-resource, *fast*-growth environments. This axis was strongly related to precipitation and moderately to temperature, with faster vegetation being found in wetter ecosystems. This finding aligns with Schellenberger Costa et al. (2017), who found leaf *acquisitive* traits to be positively associated with precipitation on Kilimanjaro. Its availability is thought to be a key driver of resource *acquisition* strategies as high availability allows plants to maintain higher rates of transpiration, photosynthesis, and nutrient uptake (Li et al., 2017). In contrast, global level analysis shows that *conservative* traits in tropical forests are related to precipitation while *acquisitive* traits relate to temperature (Bouchard et al., 2024). These contrasting results could be due to the small number of traits used by Bouchard et al (2024): SLA, SSD and seed mass. Our study included additional leaf and root traits, and so may provide a more comprehensive analysis of plant strategies and their relation to climate. Our observed relationship between land use intensity and *slow-fast* strategy axis agreed with many previous results in that leaf *acquisitive* traits were related to higher disturbance rates and fertility, as represented by the land use intensity measure (Schellenberger Costa et al., 2017, Leishman et al., 2007; Carreño-Rocabado et al., 2016, Neyret et al., 2024). The second axis, spanning from *woody* to *grassy* communities, was also driven by LUI and this may reflect the selection of shorter, grassier vegetation in more frequently disturbed environments. A similar axis was reported by Coop & Givnish, (2007), who observed a clear distinction in plant functional composition between forest and grassland in Valles Caldera, New Mexico which was mainly driven by temperature. In our study system, our LUI measure was influenced also by temperature and then drive the woody-grassy axis. While this may be because land use mostly happens at low elevations, warmer temperature conditions in these environments could create soil water deficits that hinder the growth of tall trees (Andrus et al., 2018).

The observed *woody-grassy* strategy axis corresponds to a variation in both canopy height, (similar to the *plant size* spectrum of the global economic spectrum (Díaz et al., 2016)) and in specific root length and root diameter (consistent with the fine root *collaboration* axis (Bergmann et al., 2020)) (Figure. 3). While the *woody* and *collaboration* axes have been described previously as varying independently (de la Riva et al., 2016; Weigelt et al., 2021), they both respond to factors such as soil moisture, nutrient availability (de la Riva et al., 2016; Lachaise et al., 2022), and precipitation (Ding et al., 2024). The negative influence of precipitation on both axes could explain their observed congruence in this study. The non-independence of these axes also implies that the specific effects of aboveground (*woody-grassy* strategy axis) and belowground (*collaboration* axis) factors on SOC cannot be fully disentangled in this study. Despite this limitation, our results confirm hypothesis 1, that global axes of plant strategy, typically described at species level and across global gradients, are also manifested at the community-level across the elevational gradient of Kilimanjaro.

### 4.2 Mechanisms driving AGC

We observed direct influences from precipitation and land-use intensity on AGC in these tropical mountain ecosystems. These direct positive effects of precipitation and negative influence of LUI on AGC, are consistent with other findings (Yuan et al., 2018; Stegen et al., 2011; Ensslin et al., 2015). Increased precipitation enhances water availability, which is crucial for photosynthesis and improves soil moisture and nutrient availability for plant uptake. This relationship is supported by studies showing higher nutrient availability at higher precipitation ecosystems on Kilimanjaro. Since AGC is closely related to the biomass production of woody trees, precipitation plays a significant role in resource availability, directly impacting carbon storage (de Castilho et al., 2006; Luizao et al., 2004). Keith et al., 2009; Sankaran et al., 2005; Wang et al., 2014). Second, while we did not observe a direct influence of temperature on carbon storage, unlike other studies from tropical (Raich et al., 2006; Gordon et al., 2018) and temperate regions (Liu et al., 2014; Keith et al., 2009), we did find that temperature, precipitation and LUI all indirectly influenced carbon storage through trait-mediated pathways. In communities dominated by *fast*-growing and *woody* species, we observed higher carbon storage, consistent with global findings that *acquisitive* traits contribute positively to AGC (Balvanera et al., 2006; Cavanaugh et al., 2014; Finegan et al., 2015; Shen et al., 2016). Woodiness and size-based traits are linked to aboveground carbon stocks in a self-evident manner but the association of woodiness with the *collaboration* axis may reflect an additional mechanism as high scores on the *collaboration* axis indicates dominance by species with mycorrhizal associations, which enhance nutrient and water acquisition that may enable plant productivity (Weemstra et al., 2023). Our results also accord with Conti & Díaz, (2013) who found CWM plant height to be positively associated with AGC in semi-arid forest. However, empirically, more carbon storage might be expected in communities of conservative plants because these plants have slower growth rates and longer lifespans, which leads to greater biomass accumulation and prolonged carbon retention, as it was observed by Bu et al., (2019) and Shen, et al., (2019). These discrepancies might be due to different factors including resource availability (e.g., water, nutrients) and disturbance rate. In resource-rich environments acquisitive species may store more carbon following disturbances due to rapid growth. In contrast, in resource-limited environments that experience little disturbance, conservative species that are adapted to efficient resource use may grow slowly but retain their carbon, and thus associate with high carbon storage. This aspect of disturbance may operate on Mt Kilimanjaro where forests at densely-populated lower elevations have historically experienced high degrees of logging, generally targeting the more valuable high wood density tree species, important for carbon storage (Ensslin et al., 2015). Furthermore, our positive *slow-fast* strategy axis-AGC relationship may be due to ephemeral carbon storage in fast growing plants, meaning it would not be detected in more long-term carbon monitoring. These results confirm our hypothesis 2 that functional composition, as represented here by traits related to fast growth rate and woodiness, is crucial for understanding the mechanisms governing carbon storage across broad environmental gradients. Our findings also provide support for the idea that trait-mediated effects are more important than direct climatic influences in driving AGC. This has important implications for developing strategies to optimize nature-based solutions for mitigating climate change.

### 4.3 Mechanisms driving SOC

For SOC our study indicates that the direct influence of temperature is stronger than any trait-mediated influences. We found a strong negative direct effect of temperature on SOC. These findings align with previous studies of the effect of temperature on SOC globally (Balvanera et al., 2006; Cavanaugh et al., 2014; Finegan et al., 2015; Ruiz-Jaen & Potvin, 2011; Shen et al., 2016). Decomposition and soil respiration rates increase at high temperatures, resulting in lower SOC (Hopkins et al., 2012). High temperature can also be associated with high photodegradation in litter decomposition, leading to carbon loss (Gallo et al., 2006; Hussain et al., 2023), though we would expect this mechanism is of lesser importance. Unexpectedly, we found no significant direct influence of precipitation on SOC, which apparently contrasts with other findings, although they may not have controlled for trait mediated effects (Campo & Merino, 2016; Ford et al. 2012; Kim et al., 2011; C. Wang et al., 2006; Zhang et al. 2021; Xiao et al., 2007). Additionally, contrary to what is commonly observed, there was no direct influence of LUI on SOC. This is surprising given that land use change, particularly the conversion of natural forest ecosystems to tilled agricultural ecosystems, are well-known to deplete SOC (Lal, 2005).

While we did not observe significant direct effects of precipitation and LUI on SOC, it is important to note that these factors still played a crucial role through indirect pathways. The study on the effect of land use intensity and precipitation on organic carbon at Kilimanjaro by Pabst et al. (2016), which did not consider trait mediation, found that precipitation increased SOC while LUI decreased SOC by 38%. In this study we then uncover that the mechanism of effects is likely to be mediated through the *woody-grassy* strategy axis. As mentioned above, the *woody-grassy* strategy axis probably integrated variation in both the *plant size* axis and in the *collaboration* axis (Fig. 3), which both could drive SOC. In our grassy ecosystems there may be a lower supply of aboveground litter coupled with unfavorable climate for decomposition leading to little transfer to humified SOC. At the other end of the spectrum the extensive root systems of trees in taller woody communities related to ‘*outsourcing*’ strategy may increase SOC through deposition of root exudates and the turnover of fine roots, which adds organic material to the soil as roots die and decompose, as may the slow decomposition of lignified inputs (Dijkstra et al., 2021). Increased SOC in *woody* communities can also be explained by direct acquisition of organic nitrogen by symbiotic fungi, which may bypass the decomposition of carbon compounds and allow nutrient uptake in the absence of soil carbon mineralization (Averill at al., 2014; Orwin et al., 2011). Our results align with Häger & Avalos, (2017) and Blaser et al. (2014) who showed woody communities with high wood density to be related to high SOC. Together, these results highlight the significant influence of abiotic components and substrate inputs in shaping SOC across elevational gradients.

### 4.4 Future directions

A full understanding of the linkages between functional strategies and ecosystem functioning in tropical ecosystems has been hindered by (1) a lack of trait data availability in these systems (Charles, 2018) and (2) a focus on aboveground strategies (Bardgett et al., 2014; Iversen et al., 2017). Here, we tackled these challenges by combining trait data collected in the field along a large climatic gradient, at the species level for aboveground traits and at the community-level for belowground traits. This provided a unique opportunity to comprehensively understand how they influence carbon storage. We highlighted the potential role of belowground strategies, related to the dependence on mycorrhizal symbionts, in driving belowground carbon storage. Future studies should aim to better disentangle the relative response of vegetation structure and symbiotic relationships, which were combined in our *woody-grassy* strategy axis, to climate, and their repercussions on carbon storage. This would require species-specific root trait data and could help assess the climate-mediating role of plant functional strategies on global carbon patterns. Additionally, for comprehensive prediction of the mechanism underlying carbon storage in these ecosystems data on the carbon flux at entire ecosystem both above and below ground could be of important which was not available for this study.

## 5. CONCLUSION

In this paper we have shown that (1) major axes of plant functional form and function manifested at the community level across broad elevational gradients are strongly driven by both LUI and climate and (2) that these functional strategies mediate and explain large-scale gradients in carbon storage, especially AGC. This can be attributed to the fact that both carbon storage and functional composition were strongly driven by climatic variability across Kilimanjaro. The present study presents evidence of the importance of incorporating functional traits when elucidating the complexity of ecological mechanisms determines carbon storage shifts and allocation across ecosystems. Understanding these linkages is essential for addressing the challenges posed by rapid climate change, particularly in tropical mountain regions where large amounts of carbon may be either lost or gained depending on how they are managed.

## Acknowledgements

We thank the Tanzania Commission for Science and Technology (COSTECH), the Tanzania Wildlife Research Institute (TAWIRI) and the Mount Kilimanjaro National Park (KINAPA) for granting research and access permits. We are very thankful to the German Research Foundation (DFG) for funding through the Kili project (DFG FOR 1246) and Kili-SES project (DFG FOR 5064-through grants SCHL 1934/4-1 and MA7144/3-1). Special thanks also to the Catholic Academic Exchange Service (KAAD) and Senckenberg Biodiversity and Climate research center for granting scholarship and research stay to Dickson Mauki. And final gratitude goes to University of Dodoma who granted study leave for Dickson Mauki to pursue this study project.

## Author contributions

DM, PM, MN designed the study with contributions from MS.

DM conducted data processing and analysis with contributions from MN and PM. DM, AH, DSC, NSC, JB, AE contributed data.

DM, wrote the first draft of the manuscript paper with contributions from MN, PM, MS and all authors.

## Data availability statement

Soil carbon data are available on request to Joscha N. Becker. Above ground traits data are available on TRY database dataset 698, 699 and 700 (https://www.try-db.org/de/Datasets.php), above ground carbon data are available on paper by Ensslin et al., 2015 (paper: http://dx.doi.org/10.1890/ ES14-00492.1. Data repository herbaceous: <u>10.5281/zenodo.13929135</u> shrub: <u>10.5281/zenodo.13929112</u> tree: <u>10.5281/zenodo.13929083</u>) and root traits can be accessed from Cornejo1et al., 2020 (https://doi.org/10.3389/fpls.2020.00013). Code will be available in GitHub repository when the paper is available online.

## Supplementary materials

**Fig. S1:**
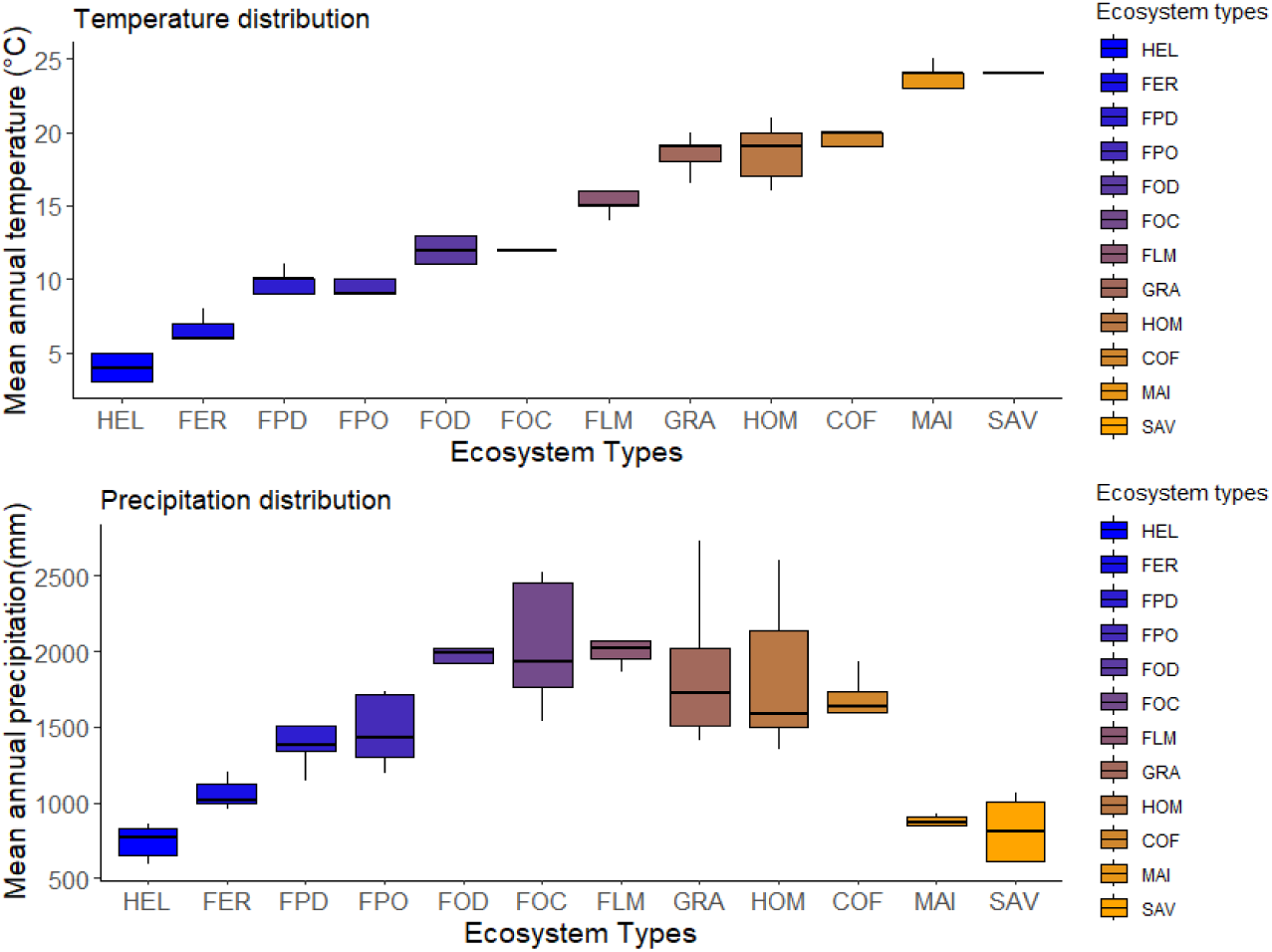
Climatic distribution of precipitation and temperature. Precipitation is low at the high elevation and lower elevation compared to middle elevation ecosystems. Mean annual temperature increased with decrease in elevation. Ecosystem types and color are arranged from high elevation to lower elevation.

**Fig. S2:**
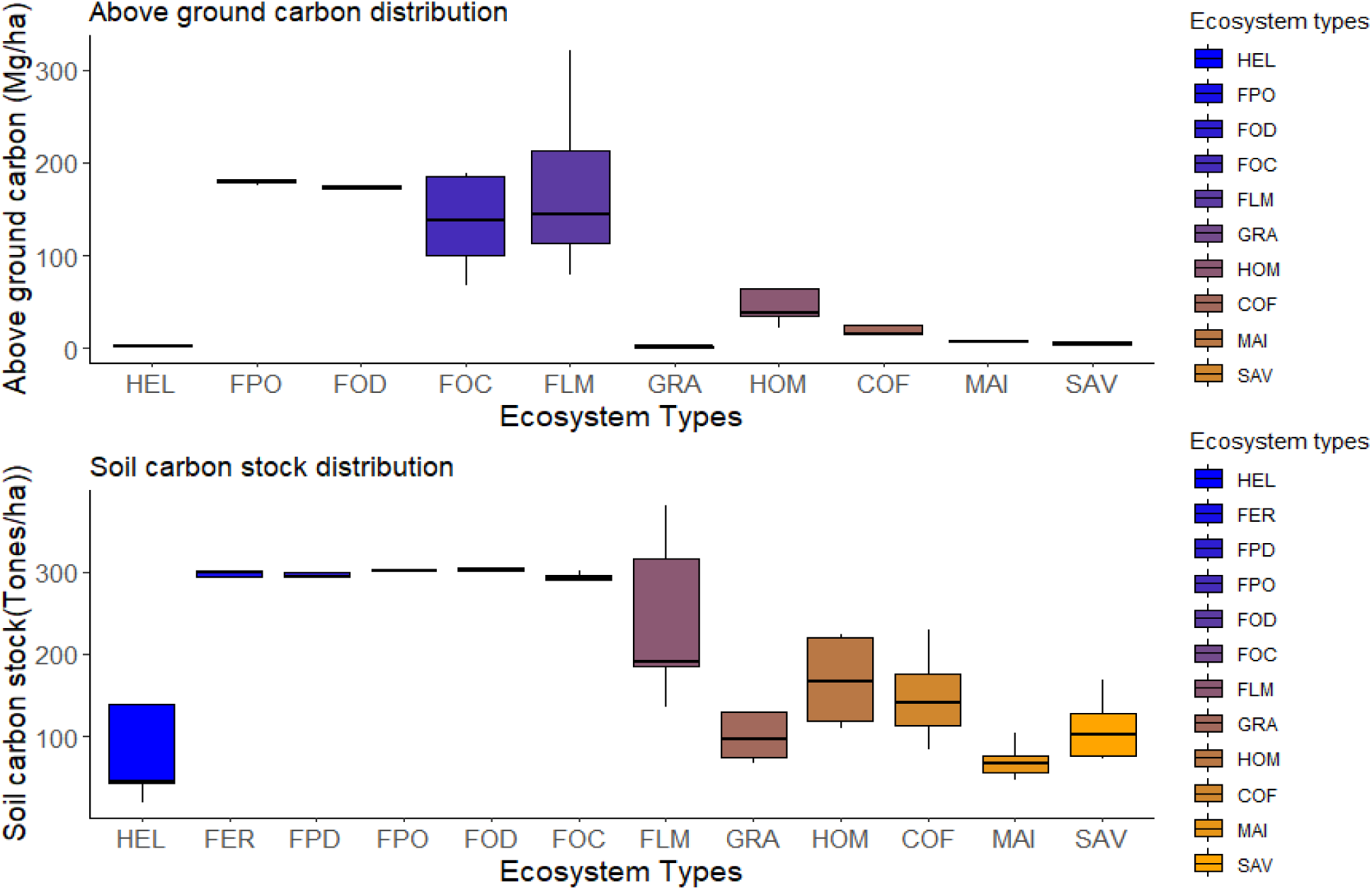
Carbon stock distribution of aboveground and soil carbon stock across elevational gradients in the Kilimanjaro ecosystem. Ecosystem types and color are arranged from high elevation to lower elevation.

**Table S1:**
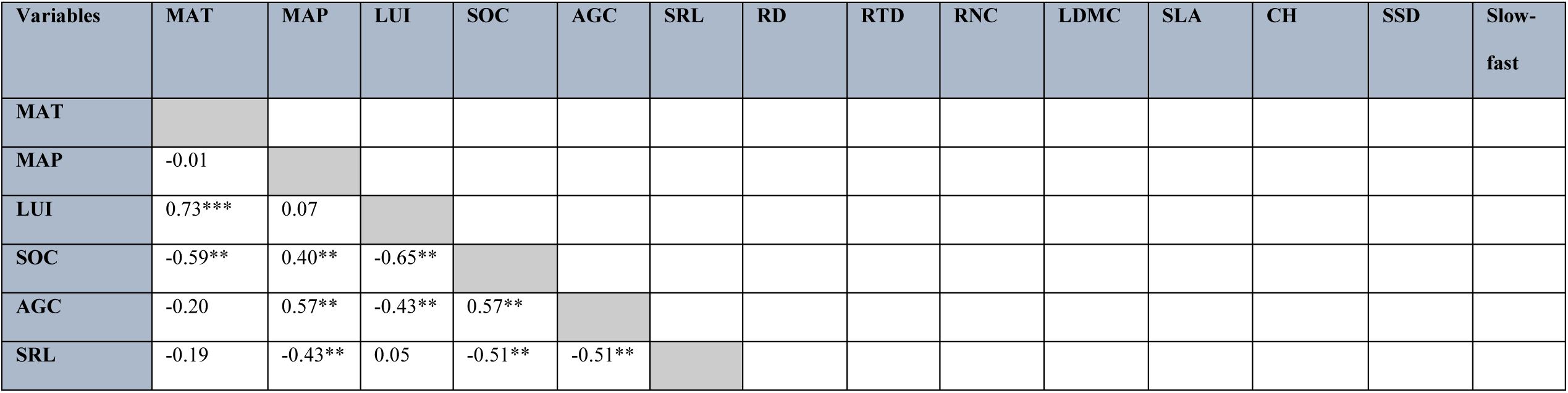
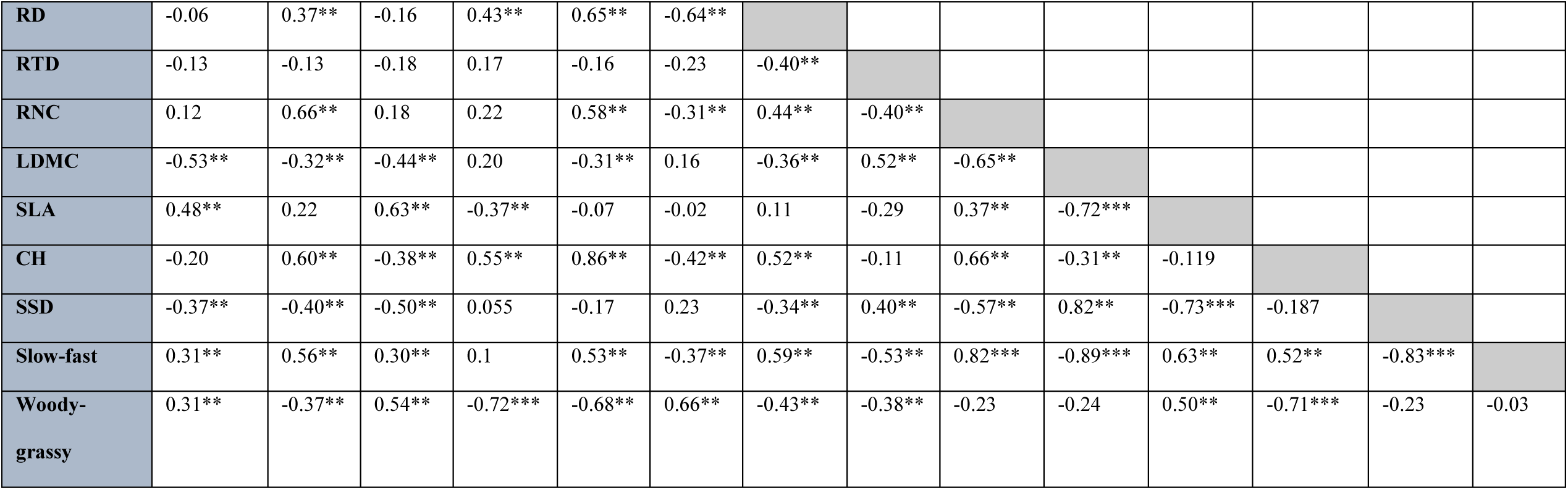
The correlation matrix table for climatic variables (MAP and MAT), LUI, functional traits and aboveground carbon and SOC and the PCA strategies. The correlation coefficient r is shown in the bottom with asterisk for significant levels (* p<0.05, ** p>0.01 and *** p>0.001).

## ALLOMETRIC EQUATIONS USED TO ESTIMATE BIOMASS ACROSS ECOSYSTEM TYPES

**Table S2.** Allometric equations used for the different habitat types and plant groups.

| Type | Allometric Equation | References | Plots |
| --- | --- | --- | --- |
| Savanna | $AGB = 0.0763 \times dbh^{2.2046} \times H^{0.4918}$ | Mugasha et al. 2013 | Savanna Maize fields |
| Wet forest | $AGB = \exp(-2.557 + 0.940 \times \ln(wd \times dbh^2 \times H))$ | Chave et al. 2005 | Low. montane forest<br>Home gardens (1, 3)<br>Coffee plantations (2)<br>Ocotea forest<br>Select. logged forest<br>Podocarpus forest |
| Dry Forest | $AGB = \exp(-2.187 + 0.916 \times \ln(wd \times dbh^2 \times H))$ | Chave et al. 2005 | Home gardens (2, 4–5)<br>Coffee plant. (1, 3–5)<br>Grasslands |
| Lianas | $AGB = \exp(-1.484 + 2.657 \times \ln(dbh))$ | Schnitzer et al. 2006 | Low. montane forest Ocotea forest Select. logged forest |
| Bananas | $AGB = 0.03 \times dbh^{2.13}$ | Hairiah et al. 2010 | Home gardens Coffee plantations |
| Coffee | $AGB = 0.281 \times dbh^{2.06}$ | Hairiah et al. 2010 | Home gardens Coffee plantations |

**Notes:** Type denotes the habitat type for which equations apply for, according to their authors.
Plots denotes the types of habitat (and for home gardens and coffee plantations also the exact plots in parentheses) for which the equations were used. Measured parameters used in the equations were total plant height (H), diameter at breast height (dbh) and wood density (wd, measured as dry weight divided by fresh volume). More explanation about the choice of the allometric equation is described in the Ensslin et al. (2015).

## LAND USE INTENSITY CALCULATION

### Calculation of the Land-Use Intensity Index (LUI)

The Land-Use Intensity Index (LUI) was calculated using four key components as described in Synthesis paper published from previous project in the same area by Peters et al., 2016.

### 1. Biomass Removal

The annual removal of plant biomass was estimated by averaging standardized measures of biomass removal at each study site. This included removal through mowing, cattle grazing, ploughing, fire events, logging, and firewood collection. Except for ploughing (measured on an ordinal scale: no ploughing, ploughing by hand, ploughing with a tractor), all other types of biomass removal were expressed as the percentage of standing biomass removed annually.

These estimates were based on repeated visits to each site (more than 20 visits per site) and cross-checked with land-use information provided by local landowners. All assessments were conducted by the same plant ecologist (A.H.), who has extensive knowledge of and over 25 years of experience with the study sites.

### 2. Agricultural Inputs

Inputs such as irrigation, fertilization, insecticides, fungicides, and herbicides were evaluated using an ordinal scale (no/very low, medium, high input). These assessments were based on local landowner information and personal observations by A.H.

### 3. Vegetation Structure

Structural characteristics were used to assess the difference between natural and managed habitats. Measurements included:

- **Canopy closure**, assessed with a canopy densitometer.
- **Maximum canopy height**, measured with a laser rangefinder.
- **Vegetation heterogeneity**, calculated as the Shannon–Wiener diversity of canopy cover values at different heights (1, 2, 4, 8, 16, 32, and 64 meters).

Measurements were taken at nine evenly distributed points within each 50 × 50-meter study site and averaged. Since natural vegetation varies along elevation gradients, raw data could not be directly used. For example, natural forests in sub montane areas typically have 100% canopy closure, whereas savanna ecosystems naturally have about 10%. To address this, the Euclidean dissimilarity of each site’s vegetation structure was calculated relative to reference sites with natural vegetation (defined to have a dissimilarity value of 0).

### 4. Landscape Composition

To capture the impact of land-use intensification at the landscape scale, the proportion of agriculturally managed habitats within a 1,500-meter radius around each site was included as a variable. This was calculated using a land-cover classification of the Kilimanjaro region, which identified 27 habitat types (18 natural and 9 managed). The classification was derived from four Terra-ASTER satellite images (11 February 2005, 2 November 2008, 28 February 2011, and 24 February 2013).

### Standardization and Composite LUI Calculation

Each component of the LUI was standardized. This ensured all components were on a uniform scale, producing LUI values ranging from 0 (lowest land use) to 1 (highest land use). The final LUI index was calculated as the average of the four standardized components.

### Validation and Alternative Approaches

The composite LUI index showed strong correlation with an index based solely on vegetation structure and landscape composition (r = 0.95, p < 0.001). Alternative measures of land use—such as using individual LUI components, a binary land-use variable, or indices with randomized weights for the components—did not alter the overall patterns observed. Furthermore, no single component outperformed the full LUI index in explaining biodiversity and ecosystem functions, underscoring its robustness.

## CLIMATIC VARIABLES MEASUREMENT AND PREDICTION

We appreciate the reviewer’s concern regarding the accuracy of the Mean Annual Precipitation (MAP) model used in our study. To ensure the robustness of our conclusions, we relied on a high-resolution precipitation dataset that has been extensively validated using independent observations and advanced geo-statistical modeling techniques as shown in Kilimanjaro paper by Appelhans et al. (2015) (https://doi.org/10.1002/joc.4552)

The precipitation model employed in this study was developed using a gradient boosting model with smooth splines, integrating rain gauge observations, satellite-derived climate data, and high-resolution elevation data. The model was validated through cross-validation techniques, with performance metrics indicating strong predictive capability. Specifically, the Root Mean Square Error (RMSE) of 429 mm and the Coefficient of Determination (R² = 0.77) suggest that the model explains 77% of the observed variance in precipitation across Kilimanjaro. These values indicate that the model performs well in predicting precipitation patterns, especially given the complexity of rainfall distribution in mountainous ecosystems.

Further validation was conducted by comparing model predictions with long-term precipitation observations from 13 independent meteorological stations on Kilimanjaro (Appelhans et al., 2015). The model successfully captured elevation-dependent precipitation patterns, including the well-documented rainfall peak at ∼2200 m a.s.l., consistent with findings from earlier studies (e.g., Hemp, 2006 (https://doi.org/10.1007/s11258-005-9049-4)). In addition, seasonal precipitation patterns aligned well with observed rainfall trends, further supporting the model’s accuracy.

Given these validation results, we are confident that the MAP estimates used in our study are robust and suitable for ecological analyses. The RMSE and R² values indicate reliable predictive accuracy, and the model’s ability to reflect known precipitation gradients across the landscape further strengthens its applicability. Therefore, while we acknowledge the importance of ensuring accurate precipitation estimates, the validation results confirm that the MAP dataset used in our study is appropriate for drawing meaningful ecological conclusions as it was also shown that our field measured data was much accurate for tropical mountains compared to Global climate databases ((Hemp and Hemp, (2024), (https://doi.org/10.1371/journal.pone.0299363)

